# Direct RNA sequencing and early evolution of SARS-CoV-2

**DOI:** 10.1101/2020.03.05.976167

**Authors:** George Taiaroa, Daniel Rawlinson, Leo Featherstone, Miranda Pitt, Leon Caly, Julian Druce, Damian Purcell, Leigh Harty, Thomas Tran, Jason Roberts, Nichollas Scott, Mike Catton, Deborah Williamson, Lachlan Coin, Sebastian Duchene

## Abstract

Fundamental aspects of SARS-CoV-2 biology remain to be described, having the potential to provide insight to the response effort for this high-priority pathogen. Here we describe the first native RNA sequence of SARS-CoV-2, detailing the coronaviral transcriptome and epitranscriptome, and share these data publicly. A data-driven inference of viral genetic features and evolutionary rate is also made. The rapid sharing of sequence information throughout the SARS-CoV-2 pandemic represents an inflection point for public health and genomic epidemiology, providing early insights into the biology and evolution of this emerging pathogen.

---

The pandemic of severe acute respiratory syndrome 2 (SARS-CoV-2), causing the disease COVID-19 and originating in Wuhan, China, has spread to more than 200 countries and territories, and has caused more than 1,000,000 cases globally [1-4]. SARS-CoV-2 is a positive-sense single-stranded RNA ((+)ssRNA) virus, belonging to the *Coronaviridae* family and *betacoronavirus* genus [5]. Related betacoronaviruses are capable of infection and ongoing transmission in mammalian and avian hosts, resulting in illness in humans such as Middle East respiratory syndrome (MERS) and the original severe acute respiratory syndrome (SARS) as examples [6-7]. Based on the limited sampling of potential reservoir species, SARS-CoV-2 has been found to be most similar to bat betacoronaviruses on a genomic level, potentially indicating that bats are a natural reservoir [5,8].

The genome sequence of SARS-CoV-2 was rapidly determined and shared on January 5^th^ of 2020, being 29,903 nucleotides in length, and annotated based on sequence similarity to other coronaviruses (GenBank: MN908947.3). As the emergence of SARS-CoV-2 has escalated, genomic analyses have played a key role in public health responses, including in the design of appropriate molecular diagnostics and supporting epidemiological efforts to track and contain the outbreak [9,10]. Taken together, publicly available sequence data suggest a recently occurring, point-source outbreak, as described in online sources [10-12].

Aspects of the response assume that the biology of SARS-CoV-2 is comparable with previously characterised coronaviruses, including the annotation of genes and the estimation of molecular evolutionary rates [11-12]. It remains highly relevant to determine these features experimentally with SARS-CoV-2-specific data, potentially revealing other insights into the biology of this emergent pathogen. To address this, here we describe (i) the first native RNA sequence of SARS-CoV-2, detailing the coronaviral transcriptome and epitranscriptome, and (ii) estimates of coronaviral evolutionary rates and related timescales, based on data available at this stage of the outbreak.

Characterised coronaviruses have some of the largest genomes among RNA viruses, and express their genetic content as a nested set of polyadenylated mRNA transcripts (Figure 1), with lengths corresponding to each encoded open reading frame (ORF). These include two large ORFs, ORF1a and ORF1ab, encoded by the complete viral genome and expressed upon cell entry. Other subgenomic mRNAs are generated through a mechanism termed discontinuous extension of minus strands, encoding structural proteins (spike protein (S), envelope protein (E), membrane protein (M) and nucleocapsid protein (N)) and accessory proteins (3a, 6, 7a, 7b, 8 and 10). The subgenomic mRNAs have a common 5′ leader sequence, near-identical to that located in the 5′-UTR of the viral genome; the transcription mechanism repositions the 5′ leader sequence upstream of ORFs, with each translation start site being located at the primary position for ribosome scanning. These ORFs, although annotated, are yet to be shown as expressed experimentally. Standard sequencing technologies are unable to produce reads representing (i) complete RNA viral genomes or (ii) subgenomic mRNAs needed to verify annotated ORFs, as these methods generate short reads and have a reliance on amplification to generate complementary DNA (cDNA) sequences.

**Figure 1.**
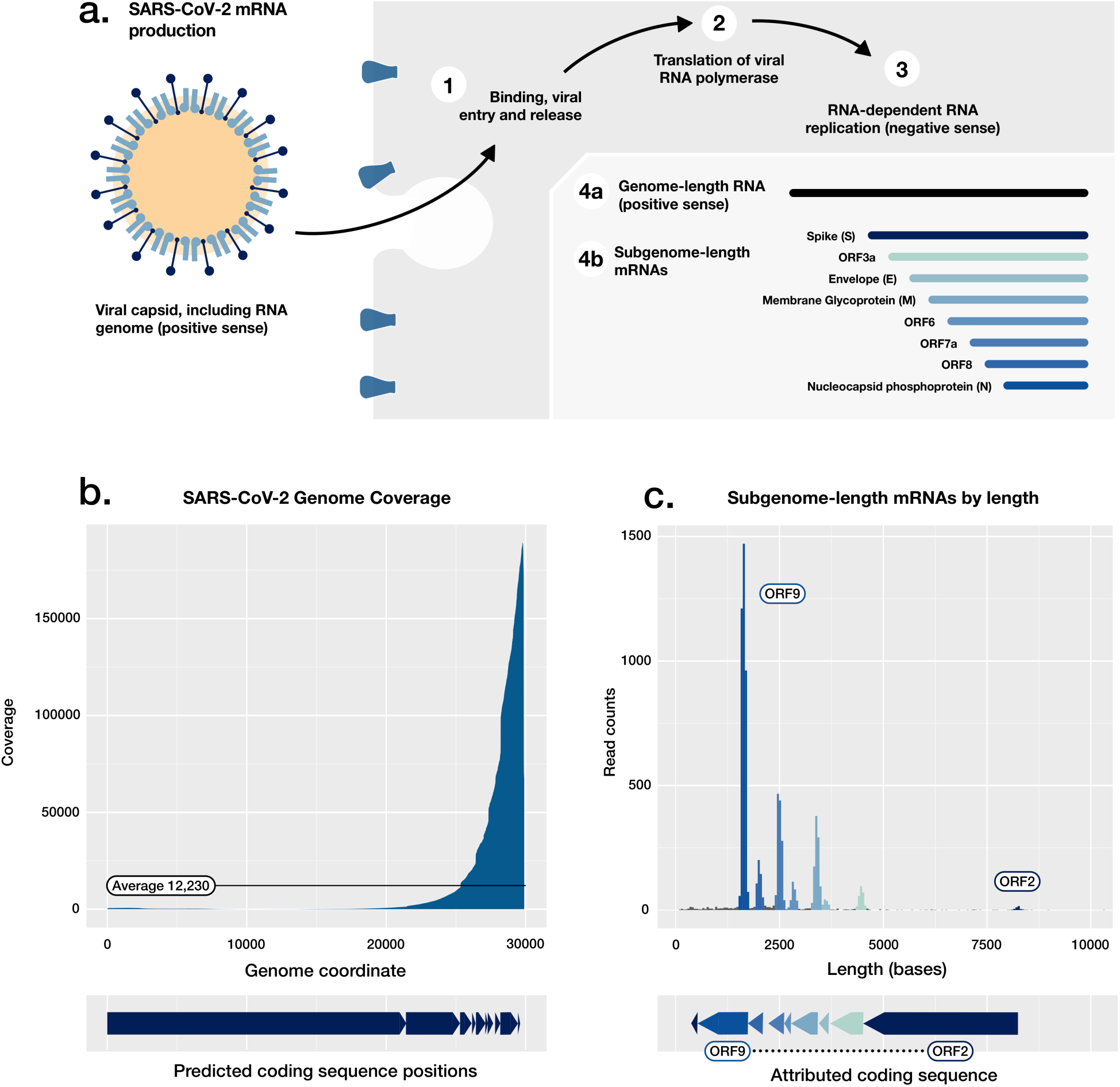
SARS-CoV-2 genetics and transcriptome architecture. A) Schematic of the early stages of SARS-CoV-2 cell entry and transcript production, including *in vivo* synthesis of positive sense genome-length RNA molecules and subgenomic mRNAs. B) Read coverage of direct RNA reads from cell-culture material, aligned to the local SARS-CoV-2 genome (29,893 bases), showing a bias towards the 3’ polyadenylated end. C) Read length histogram, showing subgenomic mRNAs attributed to coding sequences.

To define the architecture of the coronaviral transcriptome, a recently established direct RNA sequencing approach was applied, using a highly parallel array of nanopores [16]. In brief, nucleic acids were prepared from cell culture material with high levels of SARS-CoV-2 growth, this being expected to include examples of both genomic mRNA and transcripts corresponding to each ORF. These were sequenced with Oxford Nanopore Technologies, including poly(T) adaptors and an R9.4 flowcell on a GridION platform (Oxford Nanopore Technologies). Through this approach, the electronic current is measured as individual strands of RNA translocate through a nanopore, with derived signal-space data basecalled to infer the corresponding nucleobases. As a comparator, virion material of SARS-CoV-2 was also prepared and sequenced through this approach, with complete viral genome sequences expected to predominate rather than subgenomic transcripts.

The cellular-derived material was used to generate 680,347 reads, comprising 860Mb of sequence information (BioProject PRJNA608224). Aligning to the genome of the cultured SARS-CoV-2 isolate (MT007544.1), a subset of reads were attributed to coronavirus sequences (28.9%), comprising 367Mb of sequence distributed across the 29,893 base genome. Of these, a number had lengths >20,000 bases, capturing the majority of the SARS-CoV-2 genome on a single molecule. This direct RNA sequencing approach generated an average 12,230 fold coverage of the coronaviral genome, biased towards sequences proximal to the polyadenylated 3’ tail; coverage ranged from 34 fold to >160,000 fold (Figure 1B), reflecting the higher abundance of subgenomic mRNAs carrying these sequences, as well as the directional sequencing from the polyadenylated 3’ tail. The virion material generated fewer reads, and included a calibration standard added during library preparation (430,923 reads, BioProject PRJNA608224).

Many features of SARS-CoV-2 biology are captured in these direct RNA sequence data, including the transcriptome, as well as RNA base modifications or ‘epitranscriptome’. To define the transcriptome, the shared 5’ leader sequence was used as a marker to identify intact transcripts, these corresponding to subgenomic mRNAs and having a low abundance in the virion-derived data (Supplementary Figures 1 and 2). In SARS-CoV-2, we identify eight major viral mRNAs in addition to the viral genome; each annotated gene was observed as a distinct subgenomic mRNA, outside of ORF7b and ORF10 (Figure 1C, Supplementary Table 1). In SARS, ORF7a and ORF7b are encoded on a shared subgenomic mRNA, with translation of ORF7b being achieved through ribosome leaky scanning, explaining the absence of a dedicated ORF7b-encoding subgenomic mRNA [17]. There is however no satisfactory explanation for the absence of an ORF10-encoding subgenomic mRNA.

ORF10 is the last predicted coding sequence upstream of the poly-A sequence, and the shortest of the predicted coding sequences at 117 bases in length. ORF10 also has no annotated function, and the putative encoded peptide does not appear in SARS-CoV-2 proteomes [18,19] or have a homolog in the SARS-CoV-1 proteome (Proteome ID: UP000000354 [20]). Subgenomic mRNAs corresponding to ORF10 are not identifiable in our reads (Supplementary Figure 3). These data suggest that the sequence currently annotated as ORF10 does not have a protein coding function in SARS-CoV-2. Ongoing molecular evolution at this locus should be considered in light of this finding.

Instead of encoding a protein sequence, the locus annotated as ORF10 immediately upstream of the 3’ UTR may act itself or as a precursor of other RNAs in the regulation of gene expression, replication or modulating translation efficiency or cellular antiviral pathways; the 3’ UTR of coronaviruses contains domains critical for regulating viral RNA synthesis and other aspects of viral biology [21]. An initial region of the 3’ UTR appears essential for viral replication, and an area further 3’ includes the stem-loop II-like motif (s2m) a feature conserved in SARS-CoV-2 and other coronaviruses [22,23], the s2m having a proposed role in recruiting host translational machinery [24]. A small number of cell culture-derived SARS-CoV-2 genomes carry a shared deletion at an area of the 3’ UTR including an aspect of the s2m (Supplementary Figure 4), this parallel molecular evolution further suggesting the region may have functional roles *in vivo*.

An analysis of transcript breakpoints further illustrates the potential for 5’ UTR positions outside of the canonical leader sequence to enable transcript production, with low-frequency non-canonical variations in mRNA splice co-ordinates (Figure 2, Supplementary Figures 5 and 6). Low frequency variants may be generated during the preparation of nucleic acids for sequencing, with the rate of chimeric read formation being unknown; this could be explored through analysis of *in vitro* transcribed RNA control material.

**Figure 2.**
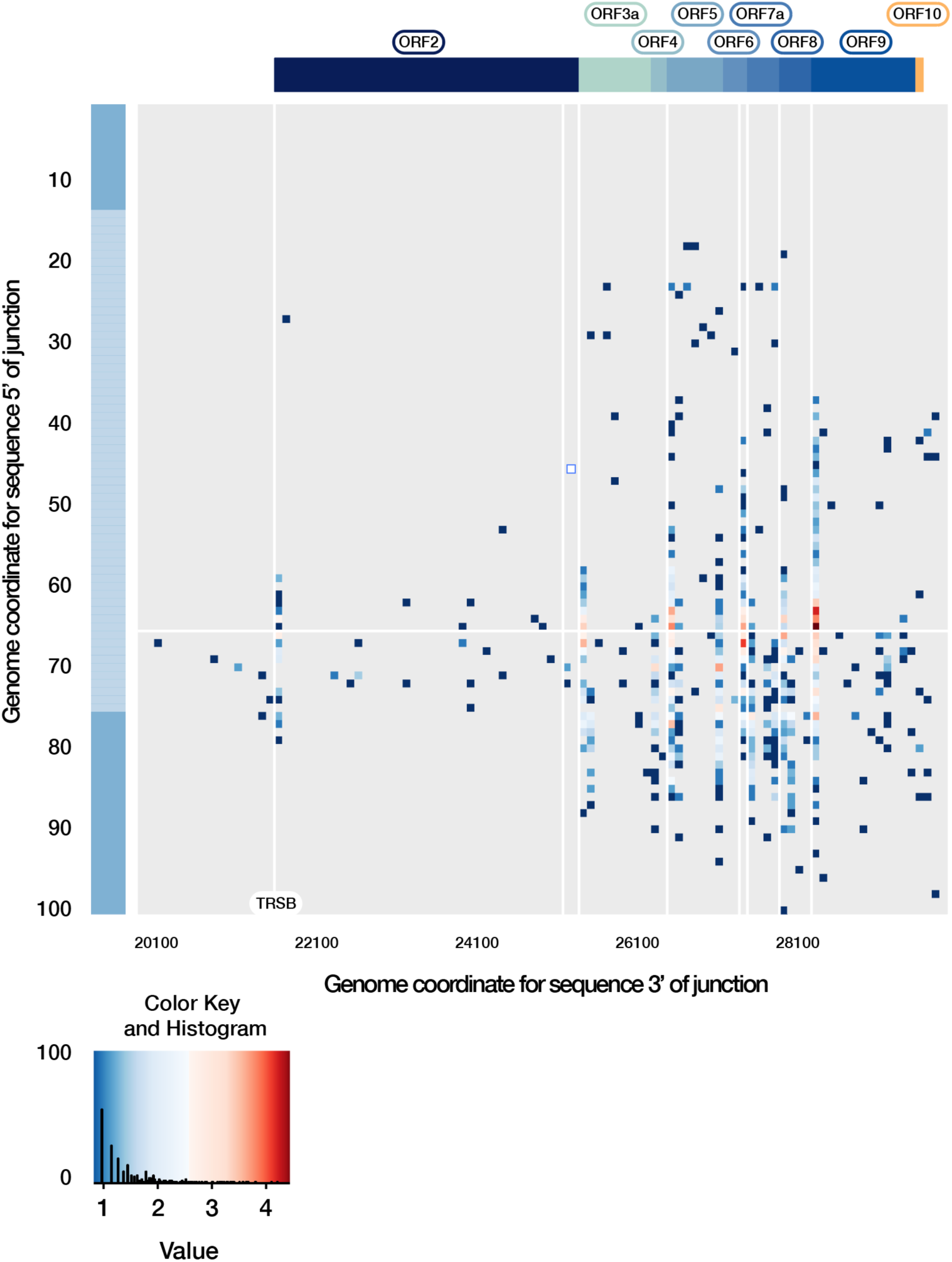
Breakpoint analysis of the SARS-CoV-2 transcriptome. Direct RNA reads carrying a breakpoint relative to the 5’ leader sequence are shown, representing potentially viable transcripts. These breakpoints are localised at the same position on the leader sequence (positions 62-68), and on the 3’ to predicted transcription regulating sequences in the body of the genome (TRS-Bs, highlighted by vertical weight lines), generating common subgenomic mRNAs. Of note, many low frequency breakpoints are detected, although few near the sequence currently annotated as ORF10. The key shows the distribution of transcript breakpoints. Colour is matched to a ‘value’ measuring the number of reads with break points at that position, log10-scaled. The histogram component illustrates the number of transcripts with a given abundance value.

In addition to RNA modifications such as the methylation of the 5’ cap structure and polyadenylation of the 3’ terminus needed for efficient translation of coding sequences, other RNA modifications may have functional roles in SARS-CoV-2 [25]. A range of modifications are identifiable using direct RNA sequence data [16,25]; our available SARS-CoV-2 direct RNA sequence data providing adequate coverage to confidently call specific modifications. Through analysis of signal-space data, we identified 42 positions with predicted 5-methylcytosine modifications, appearing at consistent positions between subgenomic mRNAs (Supplementary Figure 7, and Supplementary Table 2). In other positive ssRNA viruses, RNA methylation can change dynamically during the course of infection [26], influencing host-pathogen interaction and viral replication. Other modifications may become apparent once training datasets are available for direct RNA sequence data, with little known of the epitranscriptomic landscape of coronaviruses [25,27].

As well as investigating the above assumed features of SARS-CoV-2 genetics, sequence data also enable an estimate of the molecular evolutionary rate, with globally sourced genome sequences being shared and publicly available. Evolutionary rate estimates from other coronaviruses such as Middle East Respiratory Syndrome (MERS) are not necessarily applicable here, particularly because MERS had multiple independent introductions into humans [28-30]. To estimate the evolutionary rate and time of origin of the SARS-CoV-2 outbreak, we carried out Bayesian phylogenetic analyses using a curated set of 122 high quality publicly available SARS-CoV-2 genome sequences, each having a known collection date (Figure 3, Supplementary Table 3). The sampling times were sufficient to calibrate a molecular clock and infer the evolutionary rate and timescale of the outbreak using a Bayesian approach; the evolutionary rate of SARS-CoV-2 was estimated to be 1.20 × 10^−3^ substitutions/site/year (95% HPD 8.91×10^−4^ - 1.52×10^−3^), and the of time of origin to be late November 2019 (95% HPD August 2019, December 2019), which is in agreement with epidemiological evidence and other recent analyses (Figure 3A) [1-4, 31, 32].

**Figure 3.**
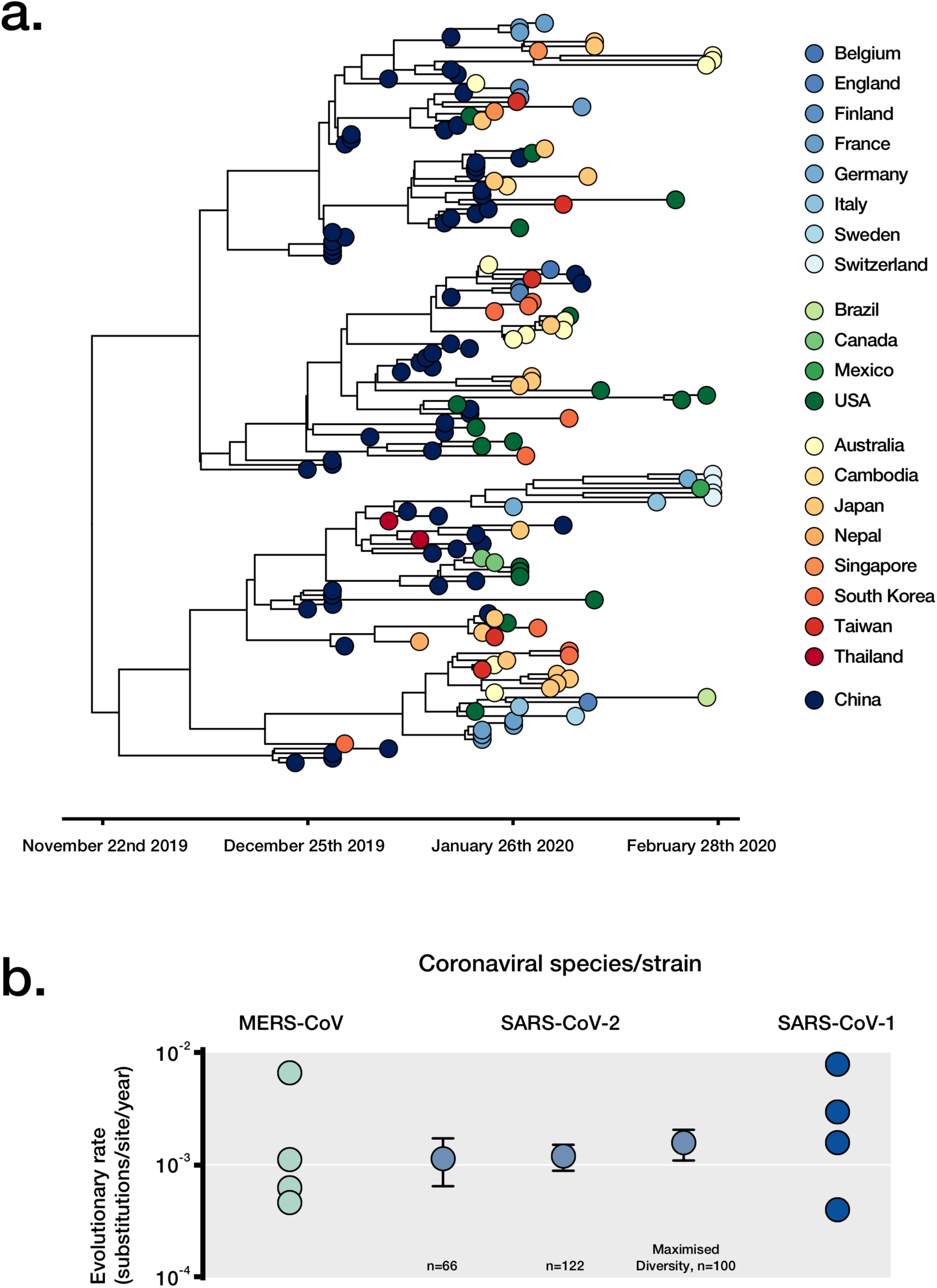
Assessment of viral evolutionary rate and outbreak timing with SARS-CoV-2-specific data. A) A timed highest clade-credibility phylogenetic tree of curated SARS-CoV-2 genomes as inferred in BEAST. B) Comparison of SARS-CoV-2 rate estimates with varying datasets, and previously published estimates of other coronaviruses.

A further set of 66 high quality genomes collected earlier in the outbreak (Supplementary Table 4), and maximal diversity data set from all data available in GISAID to March 28th to show the utility of capturing varying degrees of genetic diversity (Supplementary Table 5). This Bayesian approach demonstrated improved precision in estimates of evolutionary rates using our dataset with highest genetic diversity. This may be explained by the stochastic variation typical in data from early in an outbreak having a smaller impact as the virus accumulates genetic variation. These results are also supported by root-to-tip regression, a visual assessment of the degree of clocklike evolution in the data (Supplementary Figures 8 and 9). The evolutionary rate generated the high diversity set of 100 genomes was 1.56 × 10^−3^ substitutions/site/year (95% HPD 1.09×10^−3^ - 2.05×10^−3^), whereas that based on 66 genomes was 1.16 × 10^−3^ substitutions/site/year (95% HPD 6.32×10^−4^ - 1.69×10^−3^). Our estimate of the evolutionary rate of SARS-CoV-2 is in line with those of other coronaviruses (Figure 3B), and the low genomic diversity and recent timescale of the outbreak support a recently occurring, point-source transfer to humans.

Other phylodynamic inferences may soon become possible for SARS-CoV-2, as further genomic data becomes available and the sampling rate becomes more consistent. The current distribution of sampling times (Supplementary Figure 8) appears to be prohibitive to phylodynamic inference of the SARS-CoV-2 effective population size (N_e_, not included here). Although a required threshold of genomes to allow such phylodynamic investigation may have been crossed, the temporal spread of these isolates may differ too much to satisfy constant sampling assumptions underlying many phylodynamic skyline approaches inferring N_e_ over time. Again, as sampling continues a more consistent rate of sampling is likely to emerge, allowing such analyses.

Insights are provided on the molecular biology of SARS-CoV-2, revealed through the use of direct RNA sequence and publicly available data. The rapid sharing of these and other genetic data support the global response effort and represents an inflection point for communicable diseases and genomic epidemiology, with complete data shared openly and rapidly between academic and public health groups.

## Acknowledgements

The authors gratefully acknowledge the traditional peoples of the land on which the work was carried out, the Wurundjeri Woi-wurrung people of the Kulin nation.

The authors also acknowledge the broader staff of the Victorian Infectious Disease Reference Laboratory (VIDRL), their public health partners, and VIDRL’s major funder, the Victorian Department of Health and Human Services. The authors also thank David Matthews and Andrew Davidson of the University of Bristol for useful comments on the generated data, and Allison Hicks of Harvard T. H. Chan School of Public Health, for critical review of the manuscript. Lastly, the authors gratefully acknowledge the public health and academic groups contributing to the preparation, release and analysis of SARS-CoV-2 sequence data, including Nextstrain, GISAID and Virological.

---

**Supplementary Figure 1.**
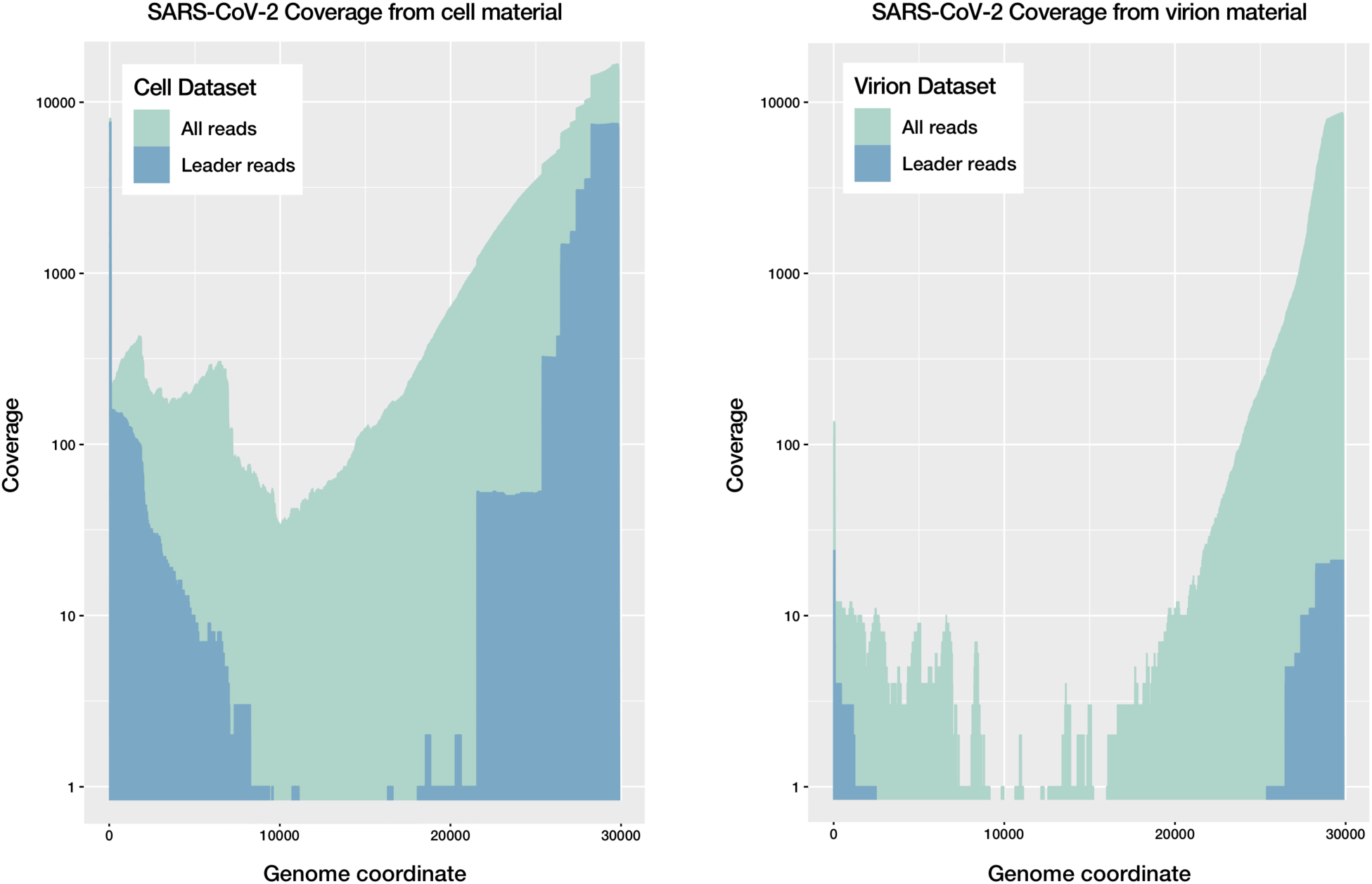
Native RNA sequence coverage of the SARS-CoV-2 genome for cell-culture and virion-derived material. A) Coverage of the SARS-CoV-2 genome for the cell-culture dataset, for all reads and for those predicted to be intact mRNA transcripts or ‘leader reads’, showing an abundance of such transcripts. B) Coverage of the SARS-CoV-2 genome for the virion-derived dataset, showing a relative paucity of intact mRNA transcripts.

**Supplementary Figure 2.**
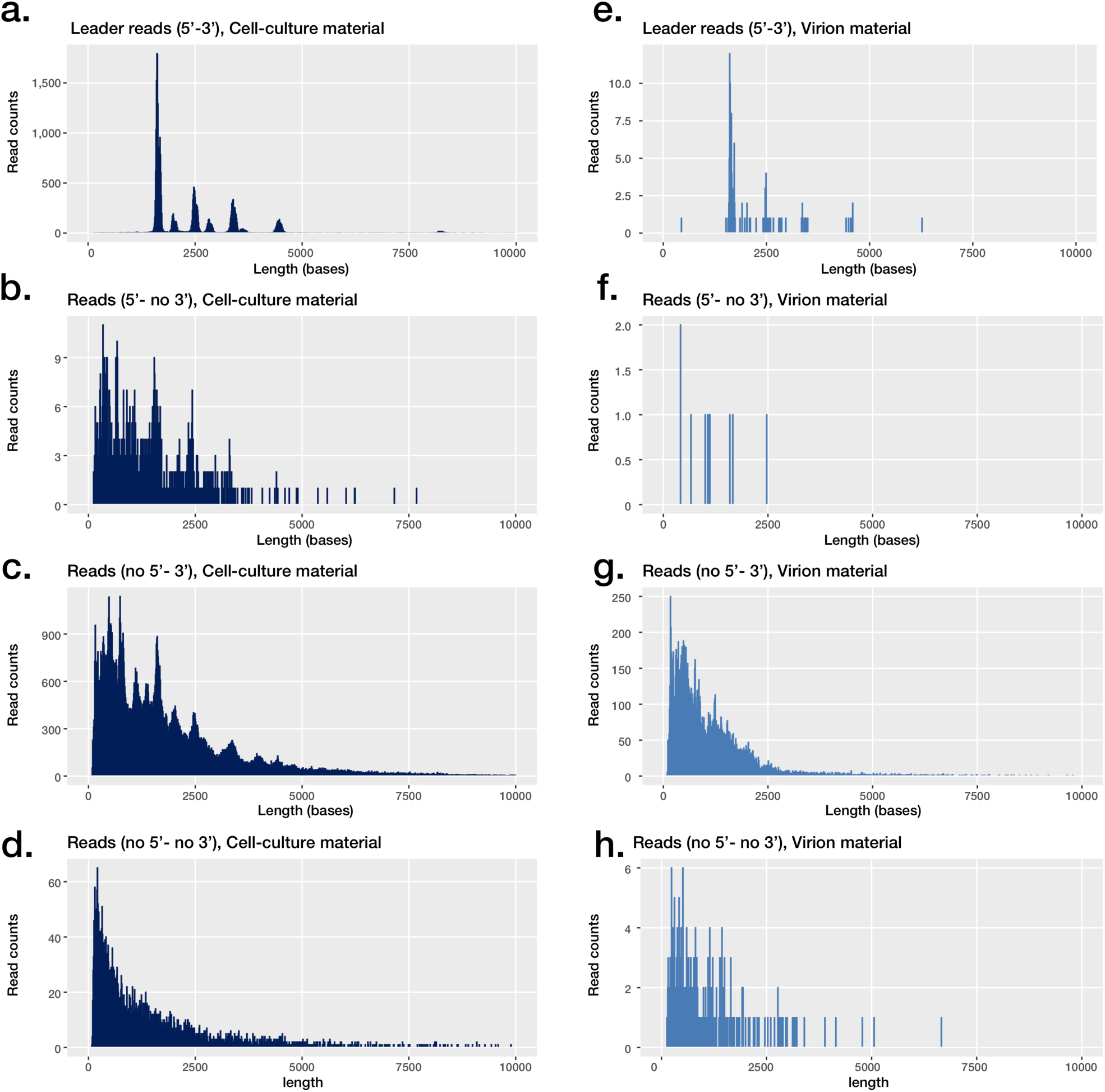
Distribution of native RNA reads between intact transcripts (‘leader reads’) and other partial transcripts and genomic sequences. Intact transcripts include the leader sequence at the 5’ and a polyadenylated 3’ end (A and E, for cell-culture material and virion material respectively), while other partial transcripts or genomes either contain a leader sequence and lack an appropriate 3’ sequence (B and F), vice versa (C and D), or lack both a leader sequence and an expected 3’ end.

**Supplementary Figure 3.**
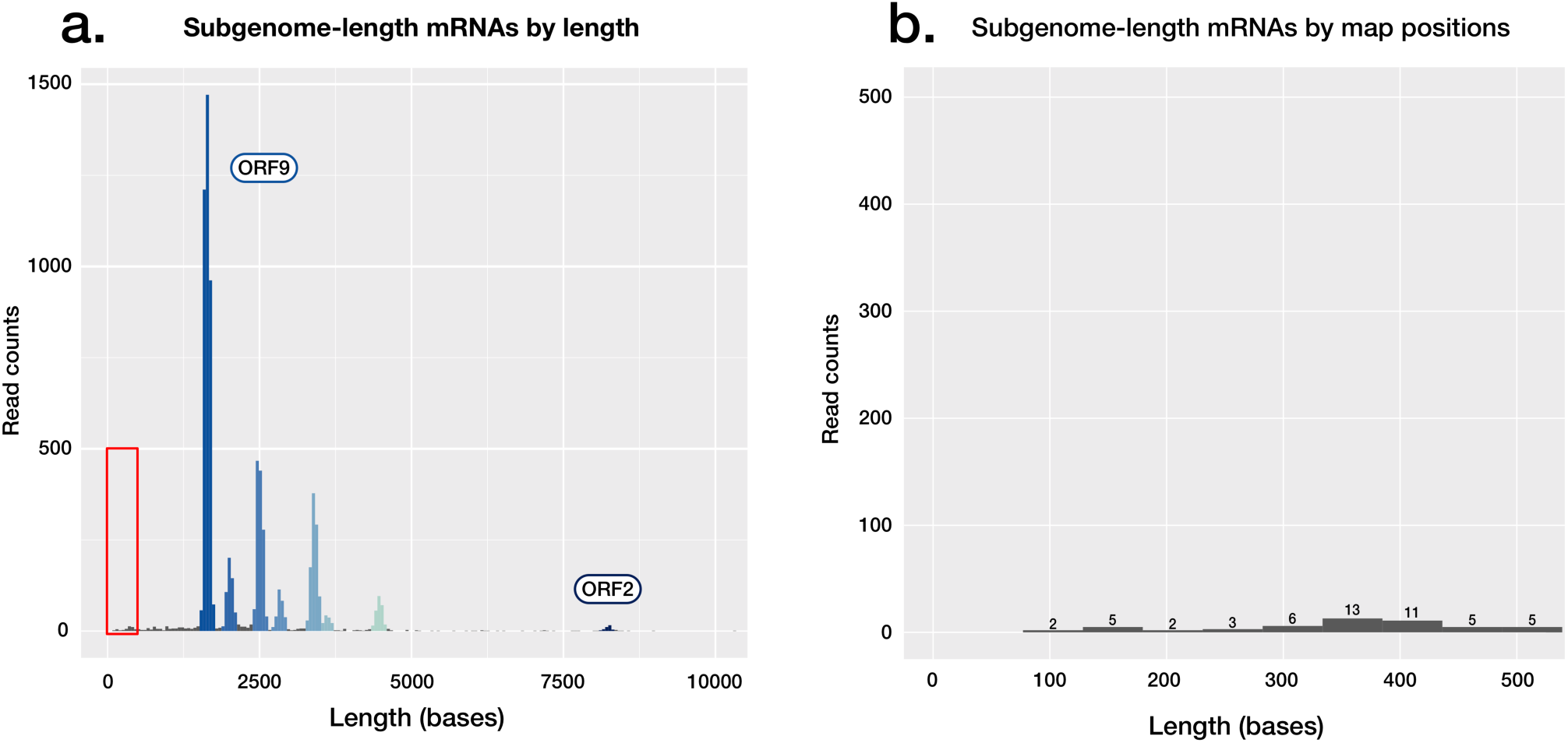
Absence of observed coding potential for ORF10 in SARS-CoV-2. A) Read length histogram, showing subgenomic mRNAs attributed to coding sequences, with the area highlighted shown in detail in a second panel. B) Read length histogram, showing read counts of lengths corresponding to those of the ORF10 subgenomic mRNA (∼360 bases), if present in the dataset. Of the <500 base reads shown, none align to ORF10.

**Supplementary Figure 4.**
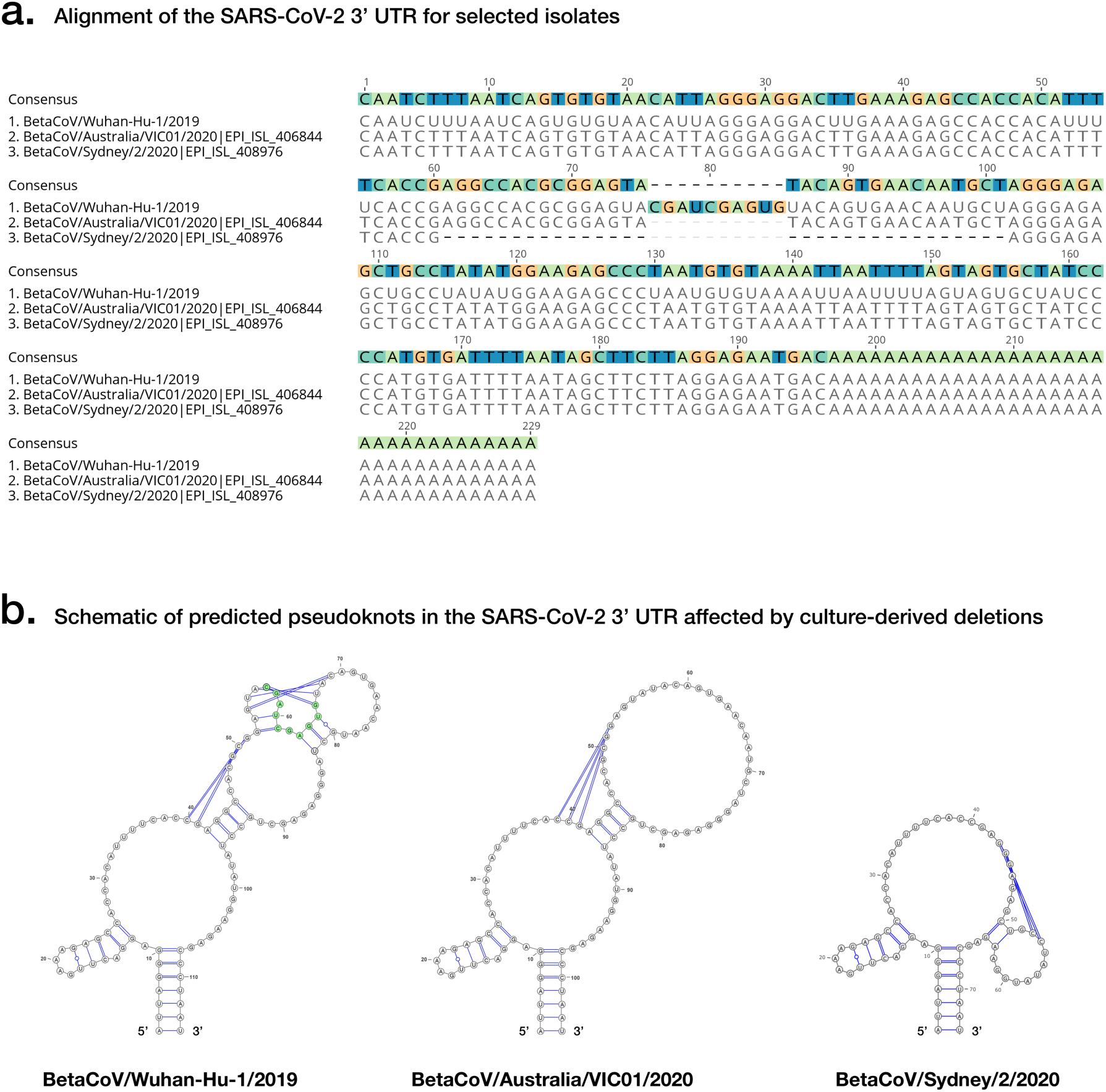
Structured RNAs in the SARS-CoV-2 3’ UTR. A) An alignment of SARS-CoV-2 3’ UTR sequences, including the original Wuhan-Hu-1 sourced from Wuhan, China and considered the reference genome for the outbreak, and two examples of cultured SARS-CoV-2 isolates exhibiting deletions in a shared 3’ UTR region predicted to form a pseudoknot structure. B) Predicted pseudoknot structure of the SARS-CoV-2 3’UTR affected by the above culture-derived deletions.

**Supplementary Figure 5.**
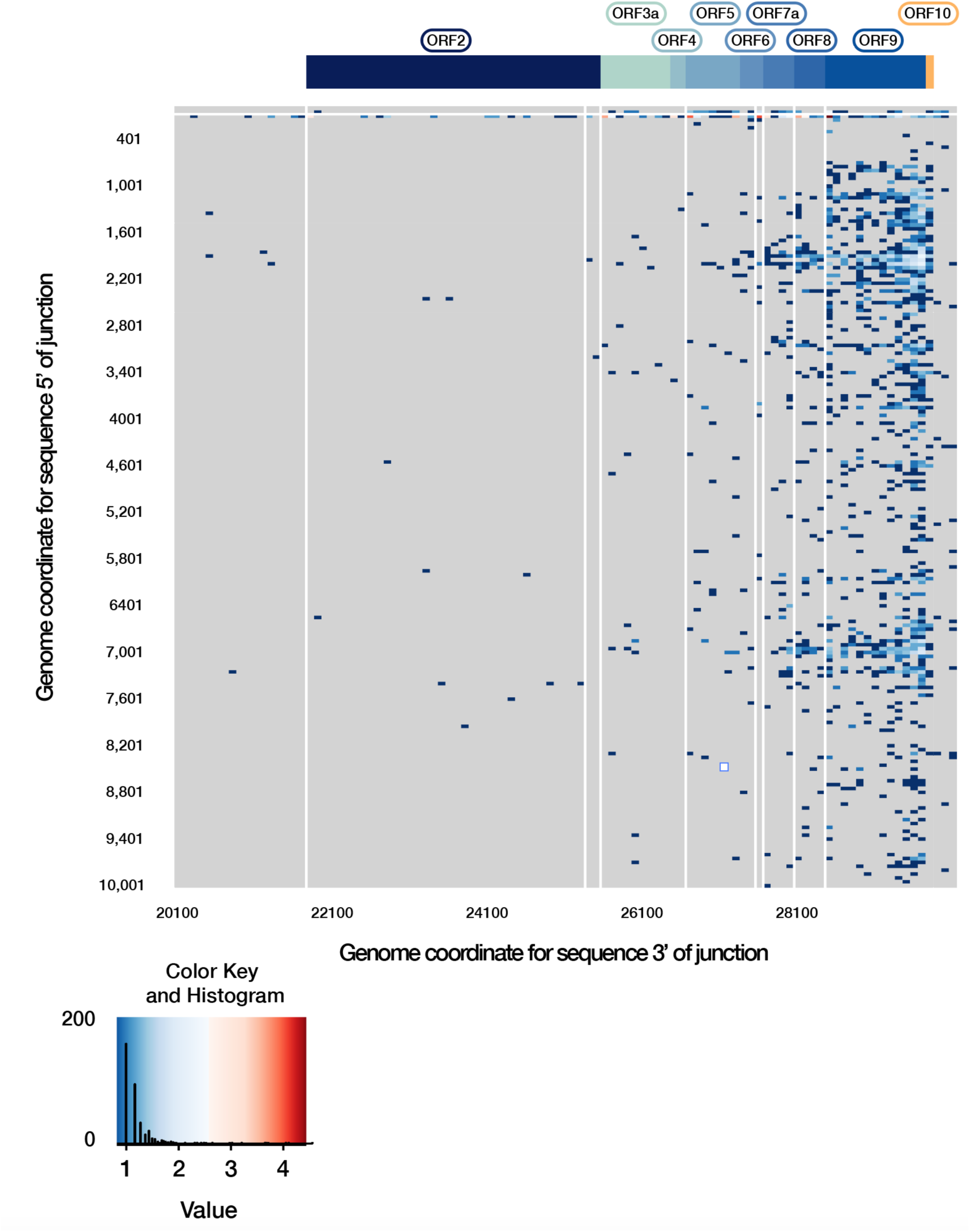
Extended breakpoint analysis of the SARS-CoV-2 transcriptome. The genome coordinates 3’ of the breakpoint are extended to include potential 3’ sequences positioned between 1-10,001 of the genome. This highlights low frequency breakpoints, increasing in frequency near the sequence annotated as ORF10 and the 3’ end of the genome. The key shows the distribution of transcript breakpoints. Colour is matched to a ‘value’ measuring the number of reads with break points at that position, log10-scaled. The histogram component illustrates the number of transcripts with a given abundance value.

**Supplementary Figure 6.**
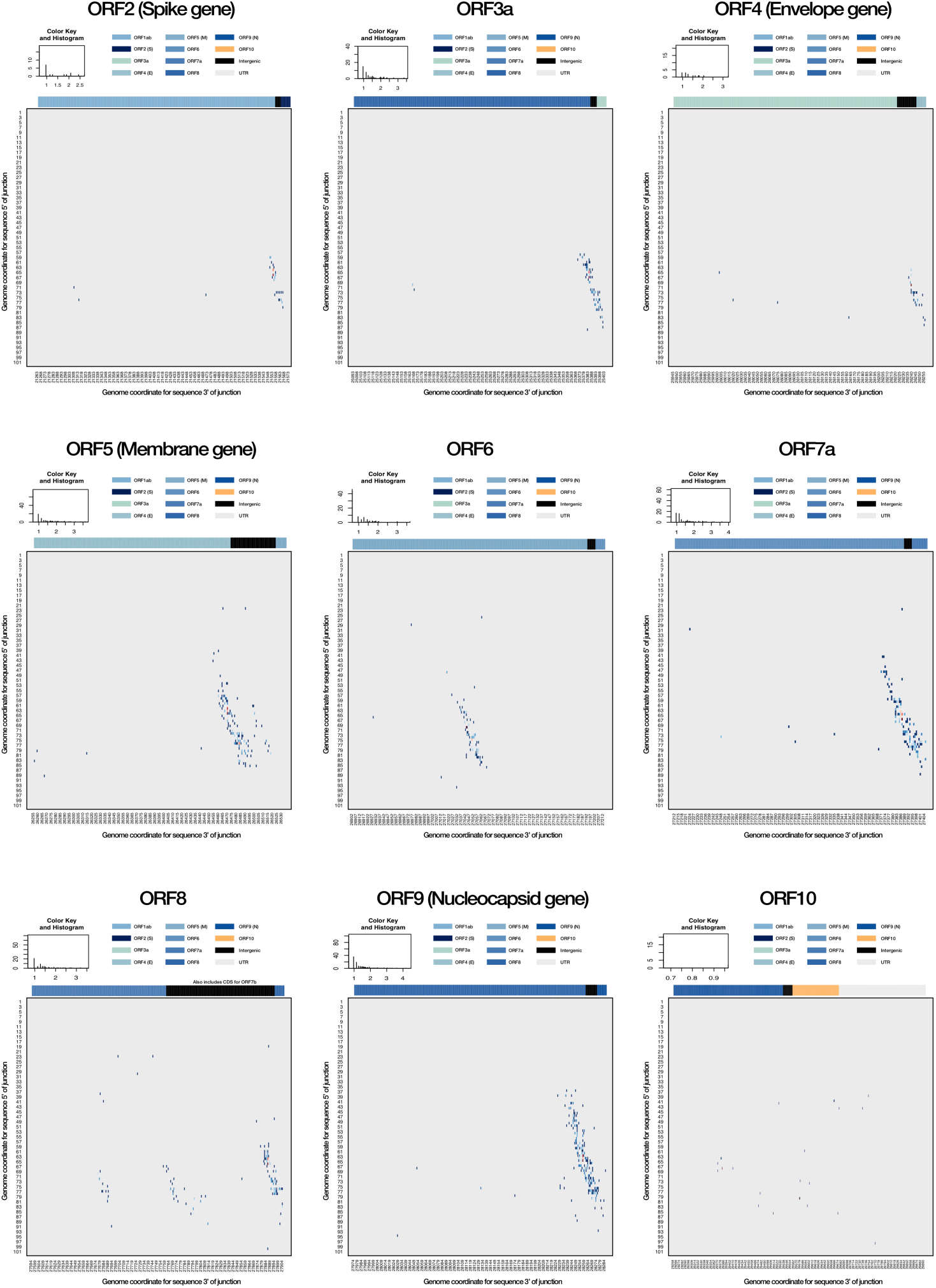
ORF-specific breakpoint analyses. The corresponding breakpoints for each currently annotated ORF in the SARS-CoV-2 genome are shown (A-I), highlighting a canonical breakpoint for ORFs with a corresponding subgenome mRNA, and a low frequency of non-canonical splice sites often centred on a canonical site. Of note, low frequency splice sites can be seen for an area between ORF 7a and ORF8, likely corresponding to ORF7b (G). There is an absence of splice sites for ORF in this dataset (I). The key shows the distribution of transcript breakpoints. Colour is matched to a ‘value’ measuring the number of reads with break points at that position, log10-scaled. The histogram component illustrates the number of transcripts with a given abundance value.

**Supplementary Figure 7.**
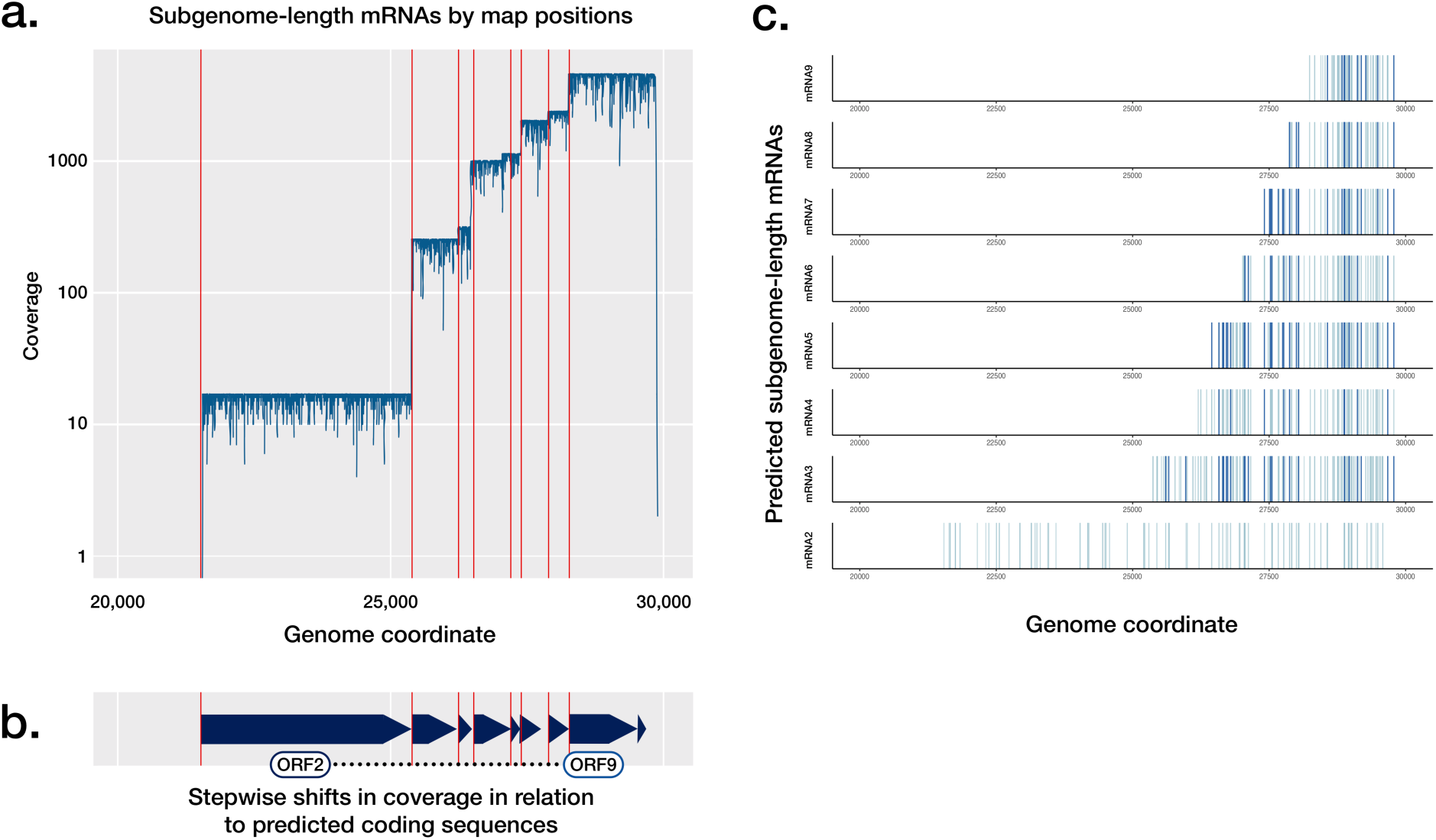
Subgenomic mRNA abundance and predicted sites of modification. A) Coverage of relevant coding sequences achieved by alignment of subgenomic mRNAs to the SARS-CoV-2 genome (log scale). Red lines indicate the first base of each coding sequence from ORF2-10. B) Schematic of relevant annotated coding sequences. C) Position of predicted m5C positions in subgenomic mRNAs. Dark blue lines indicate positions predicted to have >90% base modification; light blue lines indicate positions predicted to have between 50% and 90% base modification.

**Supplementary Figure 8.**
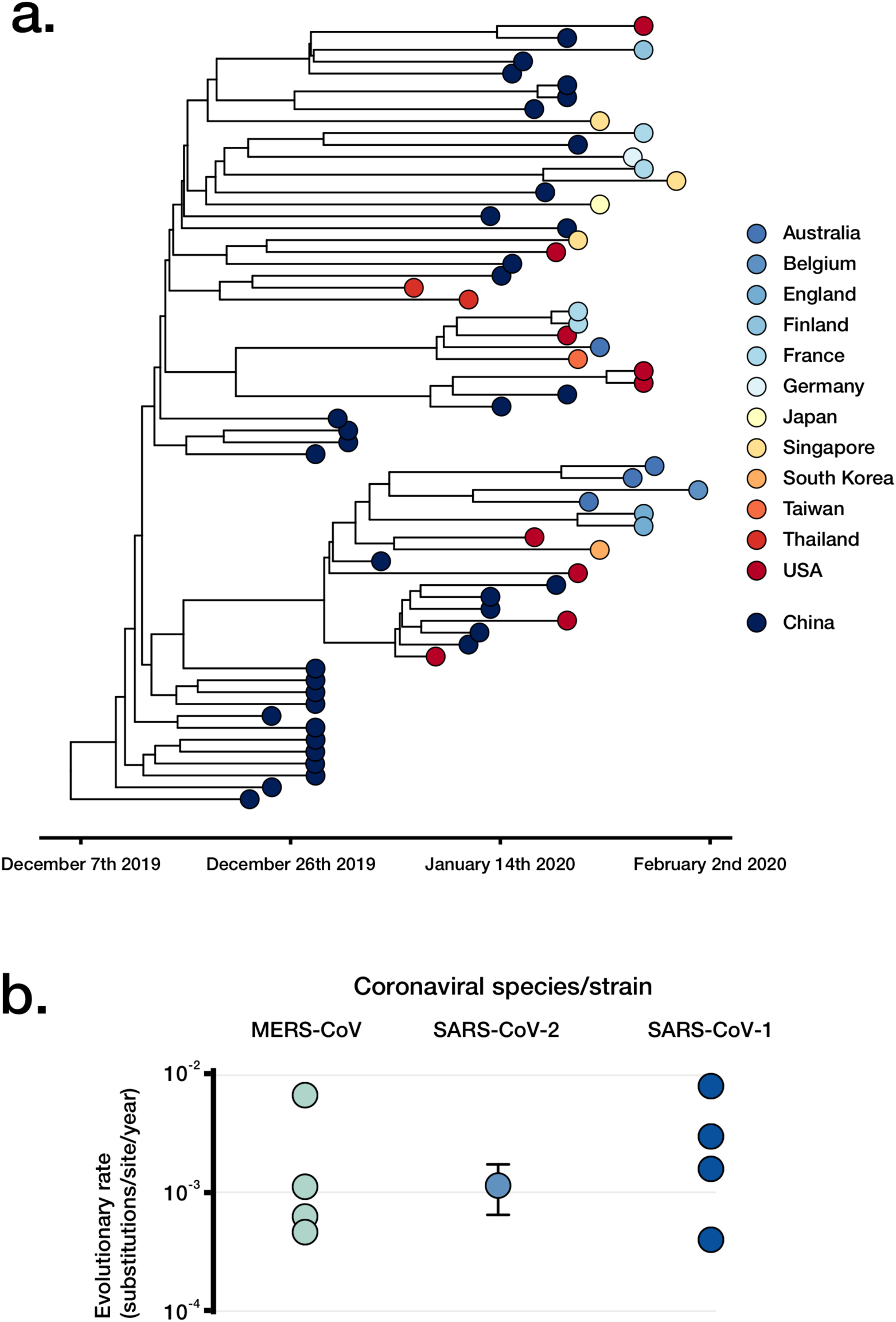
Assessment of SARS-CoV-2 phylogenetics and viral evolutionary rate based on 66 early genomes made publicly available. A) A timed highest clade-credibility phylogenetic tree of curated SARS-CoV-2 66 genomes as inferred in BEAST. B) Comparison of the SARS-CoV-2 rate estimate for the n=66 set and previously published estimates of other coronaviruses.

**Supplementary Figure 9.**
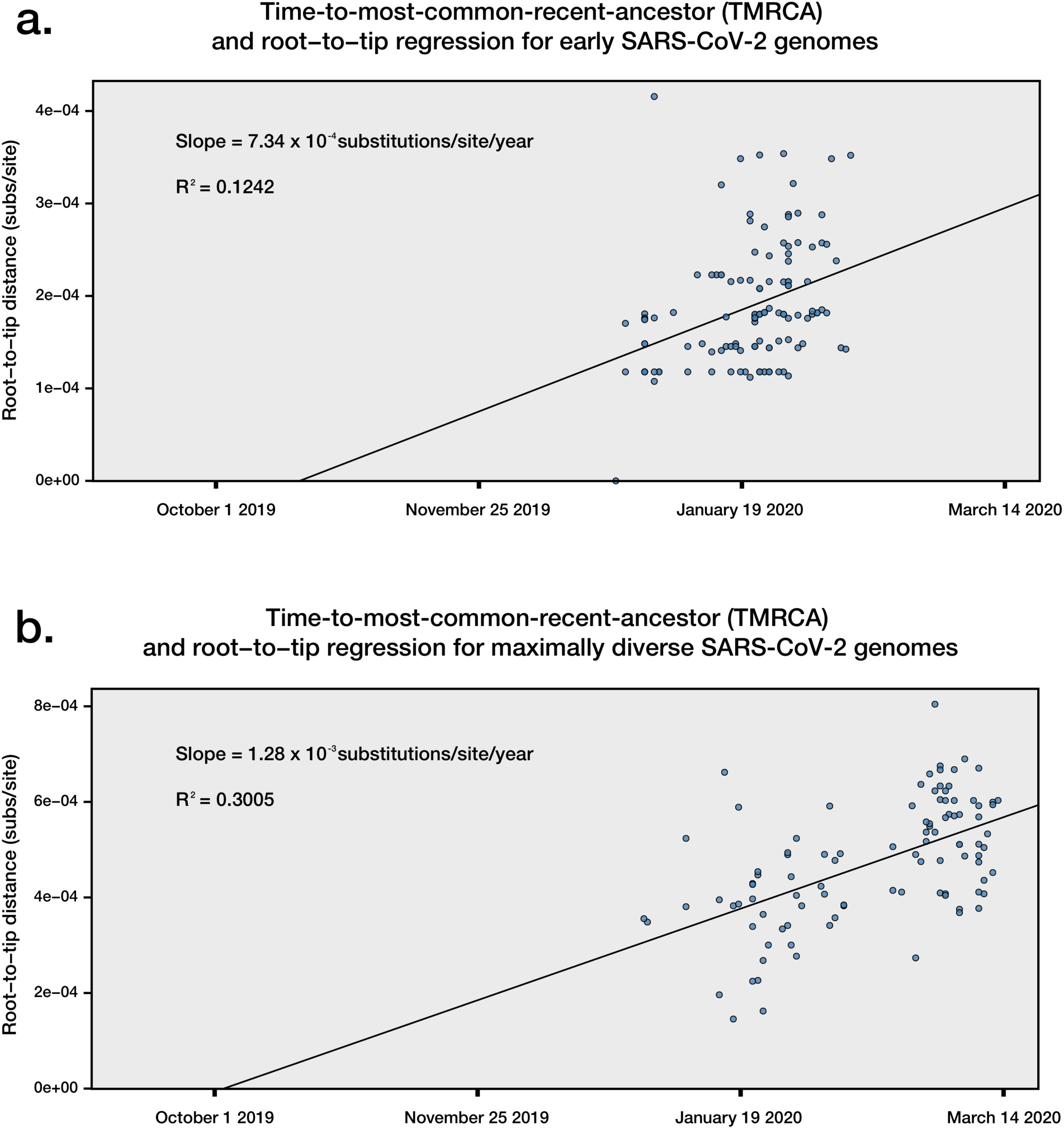
Time-to-most-common-recent-ancestor (TMRCA)and root-to-tip regression of both early and maximally diverse SARS-CoV-2 genome datasets. A) TMRCA and root-to-tip regression of 122 high quality complete SARS-CoV-2 genomes made available early in the pandemic. B) TMRCA and root-to-tip regression of 100 maximally diverse SARS-CoV-2 genomes, selected from the first 700 genomes made publicly available.

**Supplementary Figure 10.**
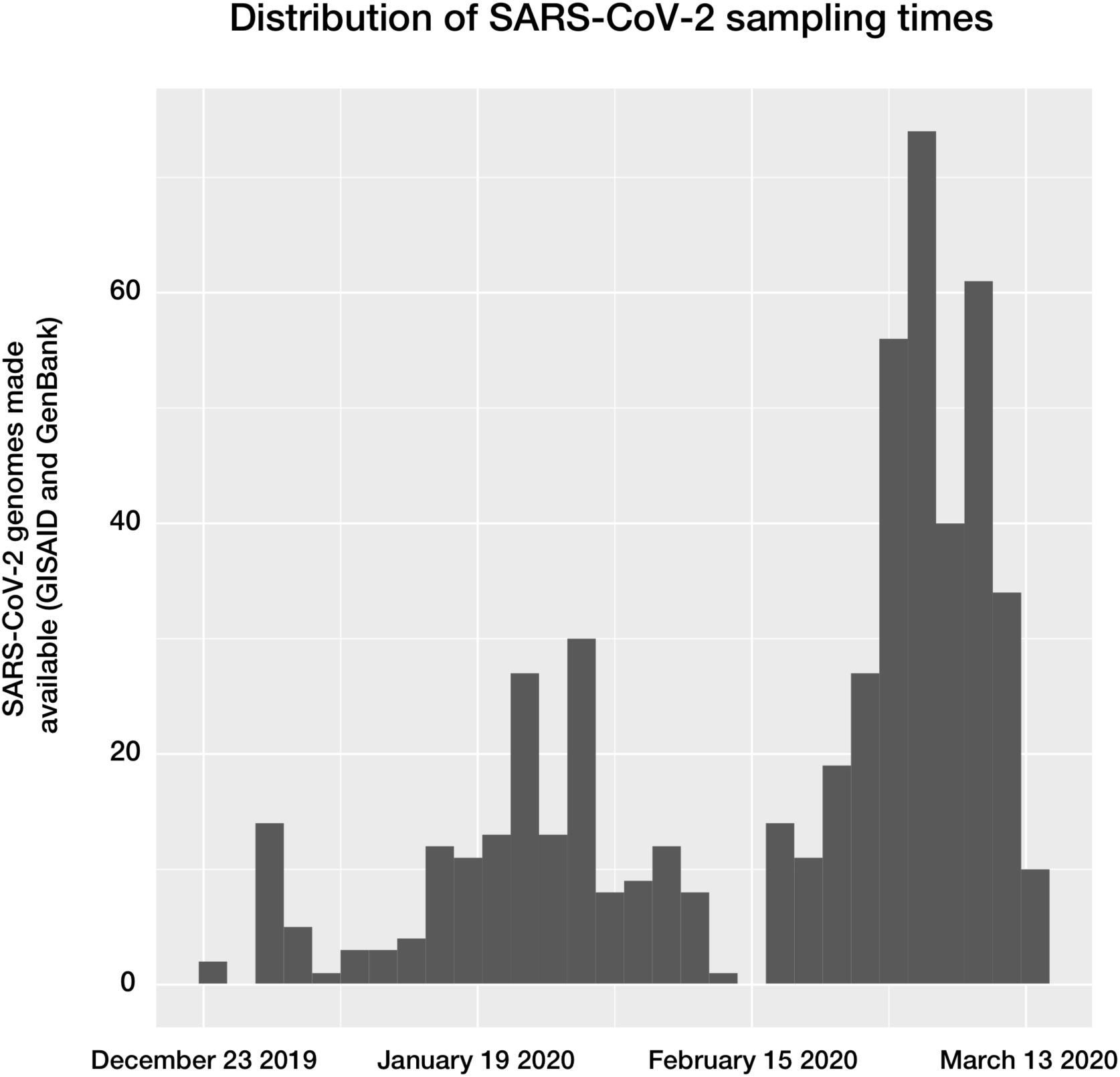
Distribution of SARS-CoV-2 sampling times used to generate publicly available genomes. The distribution has notable deviations from an expected exponential growth in the number of genomes available, such as in mid-February, with constant sampling being an underlying assumption for many phylodynamic skyline approaches inferring effective population size.

## Methods

### Samples for direct RNA sequencing

The SARS-CoV-2 material was prepared from the first Australian case of COVID-2019 (Australia/VIC01/2020), maintained in cell culture. In brief, African green monkey kidney cells expressing the human signalling lymphocytic activation molecule (SLAM; termed Vero/hSLAM cells accordingly) with associated SARS-CoV-2 infection were grown at 37°C at 5% CO_2_ in media consisting of 10 mL Earle’s minimum essential medium, 7% FBS (Bovogen Biologicals, Keilor East, Aus), 2 mM L-Glutamine, 1 mM Sodium pyruvate, 1500 mg/L sodium bicarbonate, 15 mM HEPES and 0.4 mg/ml geneticin in 25cm^2^ flasks. This isolate is to the best of our knowledge typical for SARS-CoV-2 isolates, with the genome of the cultured isolate (MT007544.1) having three single nucleotide variants (T19065C, T22303G, G26144T) relative to the SARS-CoV-2 Wuhan-Hu-1 reference genome (MN908947.3), and a 10 base deletion in the 3’ UTR. Both the T22303G and 3’ UTR variants have been confirmed as culture-derived through Sanger sequencing of clinical and culture material, and do not appear in the earlier virion-derived data.

Nucleic acids were prepared from clarified cell-free supernatant (reflecting virion material) and infected cell culture material (representing actively transcribed and translated viral material), following inactivation with linear acrylamide and ethanol. RNA was extracted from 100µl of supernatant and a modest pellet for the cell-culture material (∼200mg) respectively, using manually prepared wide-bore pipette tips and minimal steps to maintain RNA length for long read sequencing, and a QIAamp Viral RNA Mini Kit (Qiagen, Hilden, Germany). Carrier RNA was not added to Buffer AVL, with 1% linear acrylamide (Life Technologies, Carlsbad, CA, USA) added instead. Wash buffer AW1 was omitted from the purification stage, with RNA eluted in 50 μl of nuclease free water, followed by DNase treatment with Turbo DNase (Thermo Fisher Scientific, Waltham, MA, USA) 37°C for 30 min. RNA was cleaned and concentrated to 10 μl using the RNA Clean & Concentrator-5 kit (Zymo Research, Irvine, CA, USA), as per manufacturer’s instructions.

### Nanopore sequencing of direct RNA

Prepared RNA (∼1µg) was carried into a direct RNA sequence library preparation with the Oxford Nanopore DRS protocol (SQK-RNA002, Oxford Nanopore Technologies) following the manufacturer’s specifications, with addition of the control RNA in the virion sample. Libraries were loaded on R9.4 flow cells and sequenced on a GridION device for the cell-derived material and a MinION device for the virion-derived material (Oxford Nanopore Technologies), and sequenced for 40 hours. Signal-space data was used to generate nucleobase sequences (‘basecalled’) using Guppy, either as a standalone program or as ont-guppy-for-gridion 3.0.6. Both signal-space and basecalled read data are available at BioProject PRJNA608224. It should be noted that non-polyadenylated RNAs are not expected to be detected with this approach.

### Characterisation of SARS-CoV-2 transcriptome architecture

Direct RNA reads passing the above given quality thresholds were aligned to the genome of the cultured Australian SARS-COV-2 isolate (MT007544.1), with parallel and concordant analyses in Geneious Prime (2019.2.1, [M1]) and minimap2 v 2.11 using the “spliced” preset [M2]. Coverage statistics were determined from the resulting read alignments. To identify complete subgenomic mRNAs, reads were aligned to a 62 base SARS-COV-2 leader sequence (5’ACCUUCCCAGGUAACAAACCAACCAACUUUCGAUCUCUUGUAGAU CUGUUCUCUAAACGAAC), with reads aligning to the leader sequence being pooled and visualized in a length histogram. Significant peaks were identified visually and confirmed with a smoothed z-score algorithm. Reads captured in this binning-by-length strategy were re-aligned to the reference genome using the above methods and visualized in Tablet [M3]. Subgenome bins were refined to remove reads which did not originate at the 3’ poly-A tail as expected for intact subgenomic mRNAs, or which had leader sequences at least 10bp longer than expected. Subgenome bins were re-aligned, with coverage calculated in SAMtools [M4], and plotted using ggplot2 [M5] in R [M6]. Breakpoints in mRNAs were determined with CIGAR string manipulation; any given spliced region longer than 100bp (represented by Ns in the CIGAR string after aligning with minimap2) was regarded as a spliced transcript and the 5’ and 3’ genome co-ordinates of the breakpoint were recorded for analysis. The IPKnot webserver [M7] was used to predict the RNA secondary structures, and the VARNA visualization applet [M8] to produce schematics.

As an alternate method of defining the SARS-CoV-2 transcriptome, reads carrying a breakpoint relative to the 5’ leader sequence are shown, representing potentially viable transcripts. This was determined through CIGAR string manipulation. Any spliced region longer than 100bp (represented by Ns in the CIGAR string after aligning with minimap2) was regarded as a spliced transcript and the 5’ and 3’ genome co-ordinates of the breakpoint were recorded for analysis. Locations of Transcription Regulating Sequence in the body of the genome (TRS-B) were determined with a Position Weight Matrix (PWM) search. Portions of reads aligning to the conserved TRS in the leader sequence (TRS-L) were transformed into a count matrix, which was then passed into the FIMO program version 5.5.1 for motif detection with NRDB as the background distribution [M9]. Detected TRS-B sites are plotted alongside breakpoint heatmaps.

### Data availability

All signal-space (fast5) and basecalled data (fastq) generated in this work are publicly available on the sequence read archive (SRA), as part of the BioProject PRJNA608224 (See Supplementary Table 6 for relevant accession numbers).

### Identification of 5mC methylation

Nanopore sequencing preserves *in vivo* base modifications and enables their detection from raw voltage signal information. In brief, the signal-space fast5 files corresponding to identified subgenomic mRNAs were assessed to identify signal changes corresponding to 5mC methylation. These were first retrieved using the fast5_fetcher_multi function in SquiggleKit [M10]. Reads were processed to align raw signal with basecalled sequence data using Tombo v1.5 [https://github.com/nanoporetech/tombo]. Canonical reference sequences were made for each subgenomic mRNAs, with the binned fast5 files input into the detect_modifications function, with 5mC as the alternate-model parameter. Outputs were converted to dampened_fraction wiggle files and exported for visualization and analysis.

### Assessment of publicly available proteomes

Proteomic datasets were downloaded from the PRIDE proteomic database [M11] (Pride accession: PXD017710) and processes using Maxquant (1.6.3.4 [M12]) allowing semi-specific free-N-terminus tryptic as a protease specificity. Quantitation was set to TMT11 plex labelling and human proteome (Uniprot: UP000005640) and SARS-CoV2 (build in house) databases were used. Additional searches were made of the SARS-CoV reference proteome (Proteome ID: UP000000354), and recently available SARS-CoV-2 proteomic data [19].

### SARS-CoV-2 Phylogenetics

In order to estimate the evolutionary rate and time of origin of SARS-CoV-2, we carried out phylogenetic analyses in BEAST v1.101 [M13 on three datasets. The first dataset included 66 high quality genomes available up to February 10th 2020, the second consisted of 122 available up to February 24th 2020, both from GISAID and GenBank (Supplementary Table 3). A third maximal diversity dataset (n=100) was included to demonstrate the utility of capturing varying degrees of genetic diversity, these genomes being selected from the first 700 genomes available on GISAID and maximised phylogenetic diversity achieved using Treemer [M14] (Supplementary Table 4). Temporal signal was assessed using BETS [M15]. Initially we determined whether the evolutionary signal and time over which the genome data were collected was sufficient to calibrate the molecular clock, allowing for the evolutionary rate and timescale of the outbreak to be inferred. The model selection approach from BETS supported a strict molecular clock model with genome sampling times for calibration and a coalescent exponential tree prior, which posits that the number of infected individuals grows exponentially over time. We used the HKY+Γ substitution model, and set the following priors for key parameters:

- A continuous time Markov chain for the evolutionary rate
- A Laplace distribution with mean of 0 and scale of 100 for the growth rate
- An exponential distribution with mean of 1 for the effective population size.

A Markov chain Monte Carlo of length 10^7^ was set, sampling every 10^3^ steps, and assessed sufficient sampling by verifying that the effective sample size for all parameters was at least 200 as determined in Tracer [M16], automatically discarding 10% of the burn in. We summarised the posterior distribution of phylogenetic trees by selecting the highest clade credibility tree alongside calculating posterior node probabilities and the distribution of node ages. Comparison to other coronaviral evolutionary rates included studies [M17-24].

A root-to-tip regression usually produces lower evolutionary rate estimates than explicit phylogenetic methods [M25], although is commonly used to inspect temporal signal in the data. In our analyses, the data set that maximised phylogenetic diversity had a higher R^2^ than that with 122 samples collected earlier, and an evolutionary rate that was more similar to that obtained in BEAST. Although, both data sets had temporal signal according to BETS, the root-to-tip regressions demonstrate that including more genetic diversity can produce improved estimates, probably because stochastic variation has a stronger impact in smaller data sets that are collected early in the outbreak.

## References

[1] World Health Organization. Pneumonia of unknown cause — China. 2020 (https://www.who.int/csr/don/05-january-2020-pneumonia-of-unkown-cause-china/en/).

[2] United Nations Development Programme – March 2020. UNDP support for coronavirus-affected countries goes beyond health. https://www.undp.org/content/undp/en/home/blog/2020/undp-support-for-coronavirus-affected-countries-goes-beyond-heal.html

[3] Dong, Ensheng, Hongru Du, and Lauren Gardner. “An interactive web-based dashboard to track COVID-19 in real time.” The Lancet Infectious Diseases (2020). DOI: 10.1016/S1473-3099(20)30120-1

[4] World Health Organization. Statement on the second meeting of the International Health Regulations (2005) Emergency Committee regarding the outbreak of novel coronavirus (2019-nCoV). January 30, 2020 (https://www.who.int/news-room/detail/30-01-2020-statement-on-the-second-meeting-of-the-international-health-regulations-(2005)-emergency-committee-regarding-the-outbreak-of-novel-coronavirus-(2019-ncov).

[5] Lu, Roujian, et al. “Genomic characterisation and epidemiology of 2019 novel coronavirus: implications for virus origins and receptor binding.” The Lancet (2020). DOI: 10.1016/S0140-6736(20)30251-8

[6] Peiris JS, Yuen KY, Osterhaus AD, Stöhr K. The severe acute respiratory syndrome. New England Journal of Medicine. 2003 Dec 18;349(25):2431–41. DOI: 10.1056/NEJMra032498

[7] Assiri A, McGeer A, Perl TM, Price CS, Al Rabeeah AA, Cummings DA, Alabdullatif ZN, Assad M, Almulhim A, Makhdoom H, Madani H. Hospital outbreak of Middle East respiratory syndrome coronavirus. New England Journal of Medicine. 2013 Aug 1;369(5):407–16. DOI: 10.1056/NEJMoa1306742

[8] Perlman, S. Another decade, another coronavirus. New England Journal of Medicine. 2020 February 20; 382:760–762. DOI: 10.1056/NEJMe2001126

[9] Shu Y, McCauley J. GISAID: Global initiative on sharing all influenza data–from vision to reality. Eurosurveillance. 2017 Mar 30;22(13). https://www.gisaid.org/

[10] Hadfield J, Megill C, Bell SM, Huddleston J, Potter B, Callender C, Sagulenko P, Bedford T, Neher RA. Nextstrain: real-time tracking of pathogen evolution. Bioinformatics. 2018 Dec 1;34(23):4121-3. https://nextstrain.org/ncov

[11] Andersen KG, Rambaut A, Lipkin WI, Holmes EC, Garry RF. The Proximal Origin of SARS-CoV-2. Virological, accessed on 27/02/2020. http://virological.org/t/the-proximal-origin-of-sars-cov-2/398

[12] Rambaut A. Phylodynamic Analysis 129 genomes 24 Feb 2020. Virological, accessed 27/02/2020. http://virological.org/t/phylodynamic-analysis-129-genomes-24-feb-2020/356

[13] Yount B, Curtis KM, Fritz EA, Hensley LE, Jahrling PB, Prentice E, Denison MR, Geisbert TW, Baric RS. Reverse genetics with a full-length infectious cDNA of severe acute respiratory syndrome coronavirus. Proceedings of the National Academy of Sciences. 2003 Oct 28;100(22):12995–3000. DOI: 10.1073/pnas.1735582100

[14] Brian DA, Baric RS. Coronavirus genome structure and replication. InCoronavirus replication and reverse genetics 2005 (pp. 1-30). Springer, Berlin, Heidelberg.

[15] Chen Y, Cai H, Xiang N, Tien P, Ahola T, Guo D. Functional screen reveals SARS coronavirus nonstructural protein nsp14 as a novel cap N7 methyltransferase. Proceedings of the National Academy of Sciences. 2009 Mar 3;106(9):3484–9. DOI: 10.1073/pnas.0808790106

[16] Garalde DR, Snell EA, Jachimowicz D, Sipos B, Lloyd JH, Bruce M, Pantic N, Admassu T, James P, Warland A, Jordan M. Highly parallel direct RNA sequencing on an array of nanopores. Nature methods. 2018 Mar;15(3):201. DOI: 10.1038/nmeth.4577

[17] Schaecher SR, Mackenzie JM, Pekosz A (2007). The ORF7b Protein of Severe Acute Respiratory Syndrome Coronavirus (SARS-CoV) Is Expressed in Virus-Infected Cells and Incorporated into SARS-CoV Particles. Journal of Virology, 81(2), 718–731. DOI: 10.1128/JVI.01691-06

[18] Bojkova D, Klann K, Koch B, Widera M, Krause D, Ciesek S, Cinatl J, Münch C (2020) SARS-CoV-2 infected host cell proteomics reveal potential therapy targets. DOI: 10.21203/rs.3.rs-17218/v1

[19] Davidson AD, Williamson MK, Lewis S, Shoemark D, Carroll MW, Heesom K, Zambon M, Ellis J, Lewis PA, Hiscox JA, Matthews DA (2020) Characterisation of the transcriptome and proteome of SARS-CoV-2 using direct RNA sequencing and tandem mass spectrometry reveals evidence for a cell passage induced in-frame deletion in the spike glycoprotein that removes the furin-like cleavage site. DOI: 10.1101/2020.03.22.002204

[20] He R, Dobie F, Ballantine M, Leeson A, Li Y, Bastien N, Cutts T, Andonov A, Cao J, Booth TF, Plummer FA, Tyler S, Baker L, Li X (2004). Analysis of multimerization of the SARS coronavirus nucleocapsid protein. Biochemical and Biophysical Research Communications, 316(2), 476–483. DOI: 10.1016/j.bbrc.2004.02.074

[21] Goebel SJ, Hsue B, Dombrowski TF, Masters PS. Characterization of the RNA components of a putative molecular switch in the 3’ untranslated region of the murine coronavirus genome. J Virol. 2004 Jan;78(2):669–82. DOI: 10.1128/jvi.78.2.669-682.2004

[22] Tengs T, Kristoffersen AB, Bachvaroff TR, Jonassen CM. A mobile genetic element with unknown function found in distantly related viruses. Virol J. 2013 Apr 25;10:132. DOI: 10.1186/1743-422X-10-132

[23] Rangan R, Zheludev IN, Das R. RNA genome conservation and secondary structure in SARS-CoV-2 and SARS-related viruses. bioRxiv preprint 2020 DOI: 10.1101/2020.03.27.012906.

[24] Robertson MP, Igel H, Baertsch R, Haussler D, Ares M, Scott WG. The Structure of a Rigorously Conserved RNA Element within the SARS Virus Genome. PLoS Biol. 2005 Jan; 3(1): e5. DOI: 10.1371/journal.pbio.0030005

[25] Viehweger A, Krautwurst S, Lamkiewicz K, Madhugiri R, Ziebuhr J, Hölzer M, Marz M. Direct RNA nanopore sequencing of full-length coronavirus genomes provides novel insights into structural variants and enables modification analysis. Genome research. 2019 Sep 1;29(9):1545–54. DOI: 10.1101/gr.247064.118

[26] Lichinchi G, Zhao BS, Wu Y, Lu Z, Qin Y, He C, Rana TM. Dynamics of human and viral RNA methylation during Zika virus infection. Cell host & microbe. 2016 Nov 9;20(5):666–73. DOI: 10.1016/j.chom.2016.10.002

[27] Denison MR, Graham RL, Donaldson EF, Eckerle LD, Baric RS. Coronaviruses: an RNA proofreading machine regulates replication fidelity and diversity. RNA biology. 2011 Mar 1;8(2):270–9. DOI: 10.4161/rna.8.2.15013

[28] Chinese SARS Molecular Epidemiology Consortium. Molecular evolution of the SARS coronavirus during the course of the SARS epidemic in China. Science. 2004 Mar 12;303(5664):1666–9. DOI: 10.1126/science.1092002

[29] Dudas G, Carvalho LM, Rambaut A, Bedford T. MERS-CoV spillover at the camel-human interface. Elife. 2018 Jan 16;7:e31257. DOI: 10.7554/eLife.31257

[30] Cotten M, Watson SJ, Kellam P, Al-Rabeeah AA, Makhdoom HQ, Assiri A, Al-Tawfiq JA, Alhakeem RF, Madani H, AlRabiah FA, Al Hajjar S. Transmission and evolution of the Middle East respiratory syndrome coronavirus in Saudi Arabia: a descriptive genomic study. The Lancet. 2013 Dec 14;382(9909):1993–2002. DOI: 10.1016/S0140-6736(13)61887-5

[31] Andersen K. Clock and TMRCA based on 27 genomes. Virological, accessed on 27/02/2020. http://virological.org/t/clock-and-tmrca-based-on-27-genomes/347

[32] Bedford, T. Phylodynamic estimation of incidence and prevalence of novel coronavirus (nCoV) infections through time. Virological, accessed on 27/02/2020. http://virological.org/t/phylodynamic-estimation-of-incidence-and-prevalence-of-novel-coronavirus-ncov-infections-through-time/391

## References

[M1] Kearse M, Moir R, Wilson A, Stones-Havas S, Cheung M, Sturrock S, Buxton S, Cooper A, Markowitz S, Duran C, Thierer T. Geneious Basic: an integrated and extendable desktop software platform for the organization and analysis of sequence data. Bioinformatics. 2012 Jun 15;28(12):1647–9.

[M2] Li H. Minimap2: pairwise alignment for nucleotide sequences. Bioinformatics. 2018 Sep 15;34(18):3094–100.

[M3] Milne I, Bayer M, Stephen G, Cardle L, Marshall D. Tablet: visualizing next-generation sequence assemblies and mappings. Bioinformatics. 2016 (pp. 253-268). Humana Press, New York, NY.

[M4] Li H, Handsaker B, Wysoker A, Fennell T, Ruan J, Homer N, Marth G, Abecasis G, Durbin R. The sequence alignment/map format and SAMtools. Bioinformatics. 2009 Aug 15;25(16):2078–9.

[M5] Wickham H. ggplot2: elegant graphics for data analysis. Springer; 2016 Jun 8

[M6] R Core Team (2018). R: A language and environment for statistical computing. R Foundation for Statistical Computing, Vienna, Austria.

[M7] Sato, Kengo, et al. “IPknot: fast and accurate prediction of RNA secondary structures with pseudoknots using integer programming.” Bioinformatics. 2011 27.13: i85-i93.

[M8] Darty, Kévin, Alain Denise, and Yann Ponty. “VARNA: Interactive drawing and editing of the RNA secondary structure.” Bioinformatics 2009 25.15: 1974.

[M9] Grant CE, Bailey TL, Noble WS. FIMO: scanning for occurrences of a given motif. Bioinformatics 2011 27: 1017–1018

[M10] Vizcaino JA, Csordas A, del-Toro N, Dianes JA, Griss J, Lavidas I, et al. Update of the PRIDE database and its related tools. Nucleic Acids Res. 2016;44(D1):D447–56.

[M11] Cox J, Mann M. MaxQuant enables high peptide identification rates, individualized p.p.b.-range mass accuracies and proteome-wide protein quantification. Nat Biotechnol. 2008;26(12):1367–72. DOI: 10.1038/nbt.1511.

[M12] Ferguson JM, Smith MA. SquiggleKit: A toolkit for manipulating nanopore signal data. Bioinformatics. 2019 Dec 15;35(24):5372–3.

[M13] Drummond AJ, Rambaut A. BEAST: Bayesian evolutionary analysis by sampling trees. BMC evolutionary biology. 2007 Dec;7(1):214.

[M14] Menardo F, Loiseau C, Brites D, Coscolla M, Gygli SM, Rutaihwa LK, Trauner A, Beisel C, Borrell S, Gagneux S. Treemmer: a tool to reduce large phylogenetic datasets with minimal loss of diversity. BMC Bioinformatics 2018 19(1), 164. DOI: 10.1186/s12859-018-2164-8

[M15] Duchene S, Stadler T, Ho SY, Duchene DA, Dhanasekaran V, Baele G. Bayesian Evaluation of Temporal Signal in Measurably Evolving Populations. bioRxiv. 2019 Jan 1:810697.

[M16] Rambaut A, Drummond AJ, Xie D, Baele G, Suchard MA. Posterior summarization in Bayesian phylogenetics using Tracer 1.7. Systematic biology. 2018 Sep;67(5):901.

[M17] Cotten M, Watson SJ, Kellam P, Al-Rabeeah AA, Makhdoom HQ, Assiri A, Al-Tawfiq JA, Alhakeem RF, Madani H, AlRabiah FA, Al Hajjar S. Transmission and evolution of the Middle East respiratory syndrome coronavirus in Saudi Arabia: a descriptive genomic study. The Lancet. 2013 Dec 14;382(9909):1993–2002.

[M18] Cotten M, Watson SJ, Zumla AI, Makhdoom HQ, Palser AL, Ong SH, Al Rabeeah AA, Alhakeem RF, Assiri A, Al-Tawfiq JA, Albarrak A. Spread, circulation, and evolution of the Middle East respiratory syndrome coronavirus. MBio. 2014 Feb 28;5(1):e01062–13.

[M19] Dudas G, Carvalho LM, Rambaut A, Bedford T. MERS-CoV spillover at the camel-human interface. Elife. 2018 Jan 16;7:e31257.

[M20] Zhao Z, Li H, Wu X, Zhong Y, Zhang K, Zhang YP, Boerwinkle E, Fu YX. Moderate mutation rate in the SARS coronavirus genome and its implications. BMC evolutionary biology. 2004 Dec 1;4(1):21.

[M21] Salemi M, Fitch WM, Ciccozzi M, Ruiz-Alvarez MJ, Rezza G, Lewis MJ. Severe acute respiratory syndrome coronavirus sequence characteristics and evolutionary rate estimate from maximum likelihood analysis. Journal of virology. 2004 Feb 1;78(3):1602–3.

[M22] Wu SF D. CJ, Wan P, Chen TG, Li JQ, Li D, Zeng YJ, Zhu YP, He FC. The genome comparison of SARS-CoV and other coronaviruses. Yi chuan= Hereditas. 2003 Jul;25(4):373–82.

[M23] Chinese SARS Molecular Epidemiology Consortium. Molecular evolution of the SARS coronavirus during the course of the SARS epidemic in China. Science. 2004 Mar 12;303(5664):1666–9.

[M24] Sanjuán R. From molecular genetics to phylodynamics: evolutionary relevance of mutation rates across viruses. PLoS pathogens. 2012 May;8(5).

[M25] Duchêne S, Geoghegan JL, Holmes EC, Ho SY. Estimating evolutionary rates using time-structured data: a general comparison of phylogenetic methods. Bioinformatics. 2016 32(22), 3375–3379.

